# Dppa2 and Dppa4 directly regulate the Dux driven zygotic transcriptional programme

**DOI:** 10.1101/431890

**Authors:** Mélanie Eckersley-Maslin, Celia Alda-Catalinas, Marloes Blotenburg, Elisa Kreibich, Christel Krueger, Wolf Reik

## Abstract

The molecular regulation of zygotic genome activation (ZGA) in mammals remains poorly understood. Primed mouse embryonic stem cells contain a rare subset of “2C-like” cells that are epigenetically and transcriptionally similar to the two cell embryo and thus represent an ideal system for studying ZGA transcription regulation. Recently, the transcription factor Dux, expressed exclusively in the minor wave of ZGA, was described to activate many downstream ZGA transcripts. However, it remains unknown what upstream maternal factors initiate ZGA either in a Dux dependent or independent manner. Here we performed a candidate-based overexpression screen, identifying, amongst others, Developmental Pluripotency Associated 2 (Dppa2) and 4 (Dppa4) as positive regulators of 2C-like cells and ZGA transcription. In the germ line, promoter DNA demethylation coincides with upregulation of Dppa2 and Dppa4 which remain expressed until E7.5 when their promoters are remethylated. Furthermore, Dppa2 and Dppa4 are also expressed during iPSC reprogramming at the time 2C-like ZGA transcription transiently peaks. Through a combination of overexpression, knockdown, knockout and rescue experiments, together with transcriptional analyses, we show that Dppa2 and Dppa4 directly regulate the 2C-like cell population and associated transcripts, including Dux and the Zscan4 cluster. Importantly, we tease apart the molecular hierarchy in which the 2C-like transcriptional programme is initiated and stabilised. Dppa2 and Dppa4 require Dux to initiate 2C-like ZGA transcription, suggesting they act upstream by directly regulating Dux. Supporting this, ChIP-seq analysis revealed Dppa2 and Dppa4 bind to the Dux promoter and gene body and drive its expression. Zscan4c is also able to induce 2C-like cells in wild type cells, but, in contrast to Dux, can no longer do so in Dppa2/4 double knockout cells, suggesting it may act to stabilise rather than drive the transcriptional network. Our findings suggest a model in which Dppa2/4 binding to the Dux promoter leads to Dux upregulation and activation of the 2C-like transcriptional programme which is subsequently reinforced by Zscan4c.

## Introduction

Activation of transcription from the embryonic zygotic genome is a key concerted molecular and developmental event, occurring in two waves at the one to two cell stage in mouse and four to eight cell stage in humans (reviewed in (Svoboda 2017; Eckersley-Maslin et al. 2018; Jukam et al. 2017; Li et al. 2013)). Despite its importance, the precise molecular regulation of zygotic genome activation (ZGA) remains poorly understood. In particular, we still know little of the transcription factors and chromatin regulators that drive ZGA transcription, and of their coordination. Recently, the transcription factor Dux was shown to bind and activate many of such ZGA transcripts in an embryonic stem cell (ESC) model of ZGA, and be required for correct preimplantation development (De Iaco et al. 2017; Hendrickson et al. 2017; Whiddon et al. 2017). However, Dux itself is only expressed in the first or minor wave of ZGA, and what regulates Dux remains unknown.

Mouse embryonic stem cells (ESCs) represent an ideal system to study the molecular mechanism governing ZGA. Under serum or primed culture conditions, ESCs are heterogeneous and contain a small percentage of cells that not only transiently express ZGA transcripts but also share certain epigenetic characteristics with the two cell embryo (reviewed in (Eckersley-Maslin et al. 2018; Ishiuchi and Torres-Padilla 2013)). These so-called “2C-like” ESCs can be easily identified using fluorescent reporters driven by the promoters of ZGA transcripts such as the endogenous retrovirus MERVL or Zscan4 cluster (Macfarlan et al. 2012; Zalzman et al. 2010; Ishiuchi et al. 2015; Eckersley-Maslin et al. 2016). To date, repressors of the 2C-like state and ZGA gene transcription have been identified, including Kap1/Trim28 (Macfarlan et al. 2011; Rowe et al. 2010), the histone demethylase Lsd1/Kdm1a (Macfarlan et al. 2012), the histone chaperone Caf-1 (Ishiuchi et al. 2015) and the LINE1-nucleolin complex (Percharde et al. 2018) amongst others (Rodriguez-Terrones et al. 2017; Choi et al. 2017; Maksakova et al. 2013; Storm et al. 2014; Schoorlemmer et al. 2014; Hisada et al. 2012). However, aside from Dux, positive regulators that activate ZGA transcripts remain elusive.

Developmental Pluripotency Associated 2 (Dppa2) and 4 (Dppa4) are small putative DNA binding proteins expressed exclusively in preimplantation embryos, pluripotent cells and the germ line (Maldonado-Saldivia et al. 2007; Bortvin et al. 2003; Madan et al. 2009). These small proteins contain a DNA binding SAP domain and a conserved histone-binding C-terminal domain (Masaki et al. 2010; Maldonado-Saldivia et al. 2007), and physically interact and localise to euchromatin (Nakamura et al. 2011; Masaki et al. 2007). Both single and double knockout ESCs retain expression of pluripotency markers and self-renewal (Madan et al. 2009; Nakamura et al. 2011), suggesting these proteins are dispensable for stem cell pluripotency. Intriguingly, both single and double knockout mice survive early embryonic development only to develop lung and skeletal defects and perinatal lethality at a time where these genes are no longer expressed (Nakamura et al. 2011; Madan et al. 2009). This has led to suggestions that the proteins may be involved in epigenetic priming in early development, however a role in preimplantation development or in regulating ZGA transcription has not been investigated.

In order to identify new positive regulators of ZGA transcription, we performed a screen in ESCs, identifying 12 chromatin and epigenetic factors that increase the percentage of 2C-like cells within a population. Amongst these were Dppa2 and Dppa4. We investigated the regulation of these two proteins, revealing promoter DNA demethylation during the germline cycle coincides with their expression *in vivo* including in the oocyte. Knockdown of either Dppa2 or Dppa4 reduces 2C-like cells as well as expression of ZGA transcripts. Furthermore, knockout of Dppa2 and/or Dppa4 is sufficient to completely abolish this cell population. Importantly, this phenotype can be restored upon re-expression of both Dppa2 and Dppa4, but not Zscan4c, confirming these two proteins are necessary to activate expression of ZGA transcripts. Furthermore, we show that both Dppa2 and Dppa4 bind and activate Dux. Notably, Dux is required for Dppa2 and Dppa4 to activate the 2C-like state and ZGA transcription. Therefore, Dppa2 and Dppa4 act as master activators of the ZGA transcriptional programme by directly regulating the ZGA transcription factor Dux.

## Results

### Candidate-based screen for epigenetic and chromatin regulators of zygotic genome activation using 2C-like ESCs

In mouse, the major wave of zygotic genome activation (ZGA) takes place in the two-cell embryo, a stage of development which is not easily manipulated on the scale required for high-throughput screens. To circumvent this, we took advantage of a spontaneously occurring rare subpopulation of primed mouse embryonic stem cells (ESCs) that express transcripts usually restricted to ZGA, including the MERVL endogenous retrovirus and Zscan4 cluster (Macfarlan et al. 2012; Zalzman et al. 2010; Ishiuchi et al. 2015; Eckersley-Maslin et al. 2016). These “2C-like” ESCs also share several epigenetic features with the two-cell embryo, including global DNA hypomethylation (Eckersley-Maslin et al. 2016; Dan et al. 2017), decondensed chromatin (Eckersley-Maslin et al. 2016; Ishiuchi et al. 2015; Akiyama et al. 2015) and increased histone mobility (Boskovic et al. 2014).

We first performed an *in silico* screen for potential positive regulators by selecting epigenetic and chromatin regulators that are expressed in the oocyte and/or zygote (Figure 1A-B, see materials and methods). As a positive control we included Zscan4c, which has previously been implicated in activating early embryonic genes in stem cells (Hirata et al. 2012; Amano et al. 2013). Candidate genes were individually cloned as GFP fusions and transiently transfected into ESCs containing a tdTomato fluorescent reporter driven by the MERVL promoter (Eckersley-Maslin et al. 2016; Macfarlan et al. 2012) (Figure 1A). The ability of the individual genes to promote an early embryonic gene signature was tested both by flow cytometry analysis of the MERVL::tdTomato reporter (Figure 1C, D), and quantitative RT-PCR (qRT-PCR) of a panel of ZGA transcripts (Supplemental Figure 1A). Of the 22 candidates investigated, 12 promoted a 2C-like state by both flow cytometry and qRT-PCR. Following Zscan4c, *Developmental Pluripotency Associated 4* (Dppa4) was the strongest scoring screen candidate, with its closely related and interacting partner Dppa2 also amongst the screen hits. Importantly, analysis of an independent microarray dataset (Nishiyama et al. 2009) in which the transcriptome of ESCs following transcription factor overexpression was assessed, revealed that just 2 of the 50 factors investigated promoted an early embryonic transcriptome (Supplemental Figure 1B). Of the two factors that did promote expression of ZGA transcripts, Gata3 was similarly identified in our candidate-based screen, indicating that our bioinformatic preselection of candidates enriched substantially for potential ZGA regulators.

**Figure 1:**
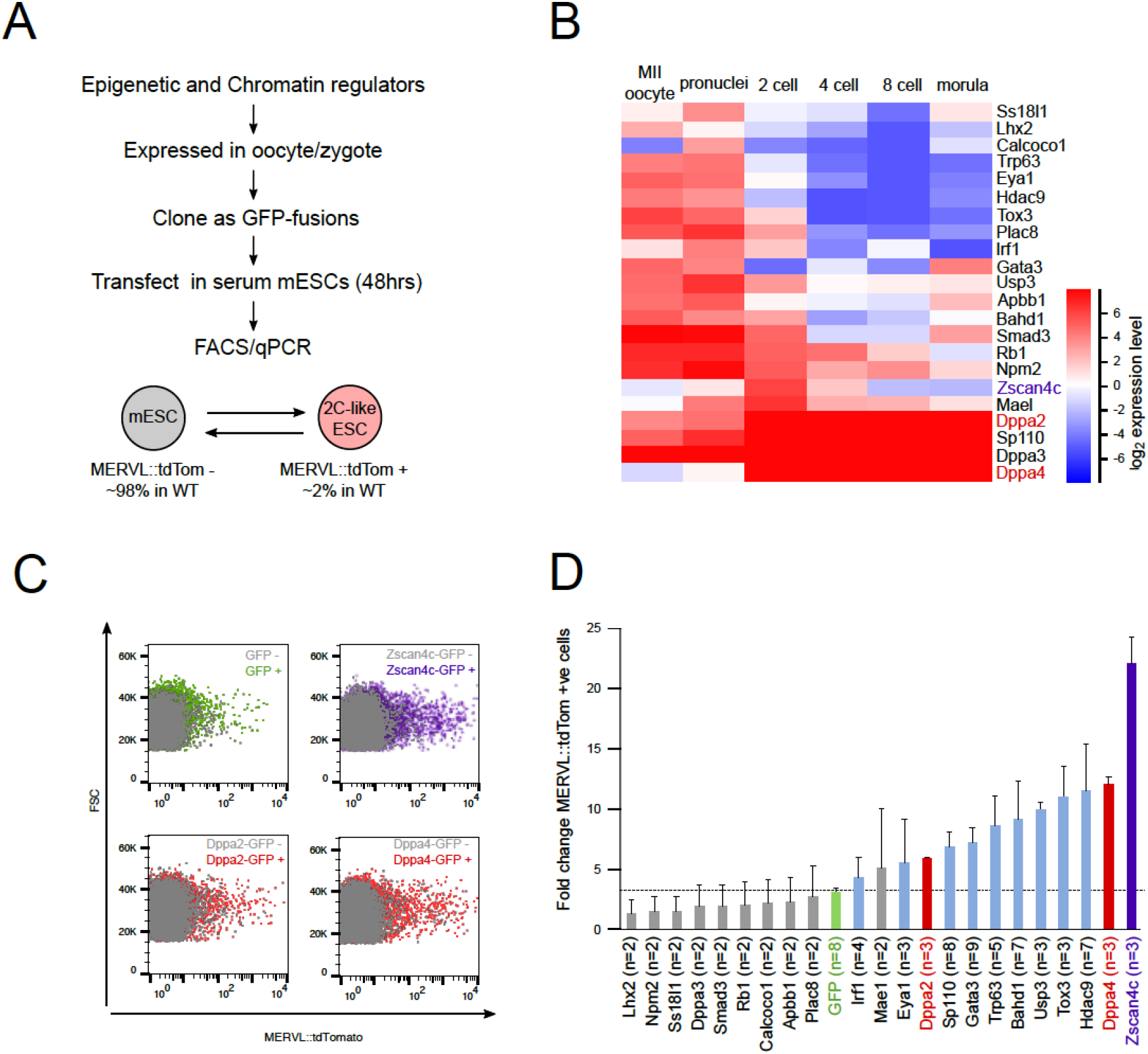
Screen for epigenetic and chromatin regulators of ZGA identifies Dppa2 and Dppa4 as potential regulators. (A) Overview of screen. Epigenetic and chromatin regulators expressed in oocyte and/or zygote were cloned and transfected in serum ESCs for 48 hours. Their ability to induce ZGA transcription was measured by flow cytometry using the MERVL::tdTomato reporter and quantitative RT-PCR on a panel of ZGA transcripts. (B) Heat map showing expression levels of factors screened in preimplantation embryos. Data from (Xue et al. 2013). (C) Representative flow cytometry plots showing levels of MERVL::tdTomato reporter (x axis) following transfection of GFP (top left), Zscan4c-GFP (top right), Dppa2-GFP (bottom left) or Dppa4-GFP (bottom right) into ESCs. Untransfected cells are shown in grey, transfected cells identified by GFP fluorescence are shown in colour indicated. (D) Fold change of MERVL::tdTomato reporter between transfected GFP positive cells over untransfected GFP negative cells following transfection of corresponding GFP-fusion constructs. GFP only control is shown in green. Bars represent average plus standard deviation of at least 2 replicates, the number of replicates is denoted for each gene.

### Dppa2 and Dppa4 activate an early zygotic transcriptional network

To validate the 12 screen hits, we performed RNA-sequencing (RNA-seq) of the GFP-positive and GFP-negative sorted cells following transient transfection of the relevant GFP-fusion construct. Transcriptome analysis confirmed an upregulation of ZGA transcripts (Eckersley-Maslin et al. 2016) in the GFP-positive sorted cells compared to GFP-negative sorted controls (Figure 2A, Supplemental Table 1 and 2). Consistently, the 12 screen hits also upregulated genes that are similarly upregulated following Dux overexpression (Hendrickson et al. 2017), CAF-1 knockdown (Ishiuchi et al. 2015) or LINE1 knockdown (Percharde et al. 2018) (Figure 2B, Supplemental Figure 2A), indicating the upregulation of 2C-like ZGA transcripts is independent of how they are defined. To accurately determine transcript levels of Dux, we remapped the RNA-seq data to a custom-built genome containing the Dux consensus sequence. Importantly, all 12 screen hits, including Zscan4c, Dppa2 and Dppa4, resulted in a significant upregulation of Dux transcript (Figure 2C). Interestingly, overexpression of Dux using a docycycline inducible transgene induced expression of several of the screen hits including the 2C-like ZGA genes Zscan4c and Sp110, as well as Dppa2 (Supplemental Figure 2B). As all three of these genes are also upregulated in 2C-like ESCs (Supplemental Figure 2C), this could suggest that positive feedback loops act to reinforce the 2C-like state.

**Figure 2:**
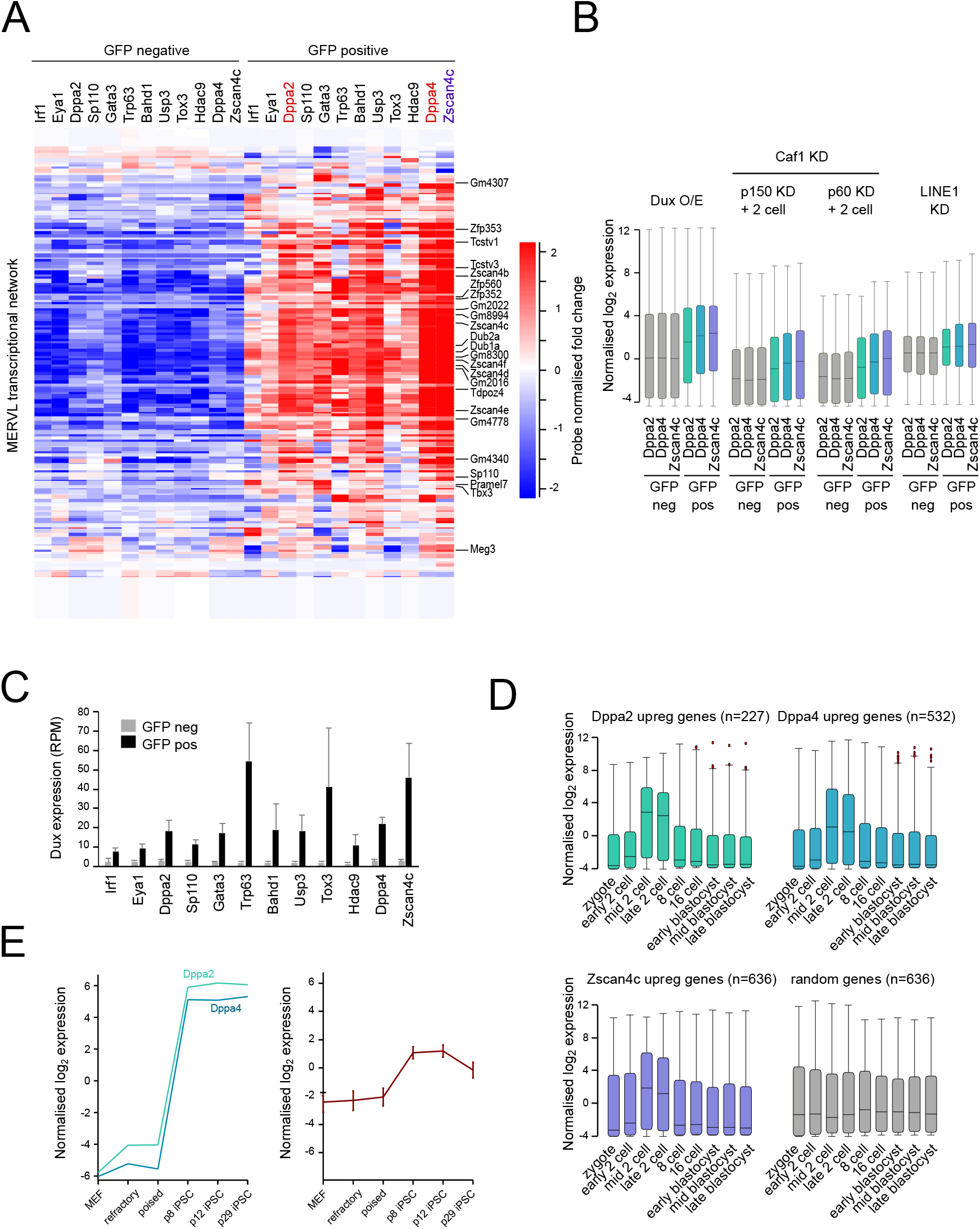
Transcriptome analysis reveals Dppa2 and Dppa4 induce transcription of ZGA genes. (A) Heat map showing per-probe normalized expression levels of ZGA transcripts expressed in 2C-like ESCs in GFP negative (left set of columns) and transfected GFP positive (right set of columns) sorted cells as measured by RNA sequencing. Gene list from (Eckersley-Maslin et al. 2016). (B) Box and Whisker plots showing expression of genes upregulated by Dux overexpression (O/E) (data reanalysed from (Hendrickson et al. 2017)), Caf1 p150 or p60 subunit knockdown (KD) and expressed in two cell embryo (gene lists from (Ishiuchi et al. 2015)) or LINE1 knockdown (gene list from (Percharde et al. 2018)) in GFP negative (grey) and GFP positive (coloured) cells following transfection of Dppa2-GFP (green), Dppa4-GFP (blue) or Zscan4c-GFP (purple). (C) Expression levels in RPM of the transcription factor Dux determined by RNA sequencing in GFP-negative sorted (grey) and GFP-positive sorted (black) cells following transfection of corresponding GFP-tagged constructs denoted below each pair of bars. Bars represent average plus standard deviation of at least 3 biological replicates. (D) Expression patterns during preimplantation development of genes upregulated by Dppa2 (green), Dppa4 (blue), Zscan4c (purple) or a random set of genes (grey). Preimplantation data from (Deng et al. 2014). (E) Expression patterns of Dppa2 and Dppa4 (left) and 2C-like ZGA transcripts (right) during iPSC reprogramming. Data reanalyzed from (Milagre et al. 2017). MEF, mouse embryonic fibroblasts. Refractory (SSEA1-/Thy1+) and poised (SSEA1+/Thy1) stages correspond to fluorescence-activated cell sorting (FACS)-sorted cells at day 6, where passage 8 (p8; corresponding to day 21), p12 (corresponding to day 29) iPSCs represent intermediate-late stages of reprogramming and p29 (corresponding to day 60) iPSCs are fully reprogrammed.

The transcriptome upregulated by Zscan4c, Dppa2 and Dppa4 showed a high degree of overlap (Supplemental Figure 2D), again indicating that a similar transcriptional network is activated following overexpression of these three factors. In all cases, the transcripts upregulated by Dppa2, Dppa4, Zscan4c or other screen hits were similarly upregulated in the mid-to-late two-cell stage during embryogenesis (Figure 2D, Supplemental Figure 2E) confirming these transcripts are activated during ZGA. To further support their role in promoting expression of 2C-like ESCs, we looked at expression patterns of Dppa2, Dppa4 and Zscan4c during iPSC reprogramming. Upregulation of 2C-like ZGA transcripts at intermediate stages of iPSC reprogramming has been previously reported (Eckersley-Maslin et al. 2016; Zhao et al. 2018). Consistently, both Dppa2 and Dppa4 are expressed when the ZGA transcripts are upregulated (Figure 2E). In summary, Dppa2 and Dppa4 upregulate an early embryonic transcriptional programme.

### Early embryonic and germline expression of Dppa2 and Dppa4 is regulated by promoter DNA demethylation

Given the specific and restricted expression pattern of Dppa2 and Dppa4 in the germ line and early embryo we investigated the regulation of Dppa2 and Dppa4 *in vivo*. Primordial germ cells (PGCs) undergo a wave of DNA demethylation which is then re-established in the mature gametes before a second wave of DNA demethylation takes place after fertilisation in the preimplantation embryo (reviewed in (Lee et al. 2014; Eckersley-Maslin et al. 2018)). Consistently, in both male and female PGCs, the *Dppa2/4* locus is demethylated (Figure 3A, Supplemental Figure 3A, B), which coincides with their expression in the gonads (Maldonado-Saldivia et al. 2007) and developing oocytes (Figure 3B). In sperm and oocytes, there is a gain in DNA methylation across the locus, however the promoter of both *Dppa2* and *Dppa4* remain hypomethylated (Figure 3A, Supplemental Figure 3A). Following fertilization, there is a second wave of DNA demethylation across the entire *Dppa2/4* locus (Figure 3A). After implantation, levels of DNA methylation, including at the promoter, increase dramatically, consistent with the rapid silencing of Dppa2 and Dppa4 (Figure 3C). The promoters of *Dppa2* and *Dppa4* remain methylated across all somatic tissues where Dppa2 and Dppa4 are not expressed (Supplemental Figure 3C). To further investigate the link between promoter DNA methylation and Dppa2/4 expression, we investigated transcriptome data from E8.5 embryos that lacked the *de novo* DNA methyltransferase Dnmt3b which is primarily responsible establishing DNA methylation at promoter regions (Auclair et al. 2014). Importantly, there was an increase in both Dppa2 and Dppa4 expression in Dnmt3b^-/-^ embryos at a time when they are usually completely silenced (Figure 3D), supporting a role for promoter DNA methylation in repressing these two genes *in vivo*. Hence the *Dppa2* and *4* genes are primarily regulated by global demethylation during germline and early embryo development, and their products are therefore present in the oocyte at fertilization.

**Figure 3:**
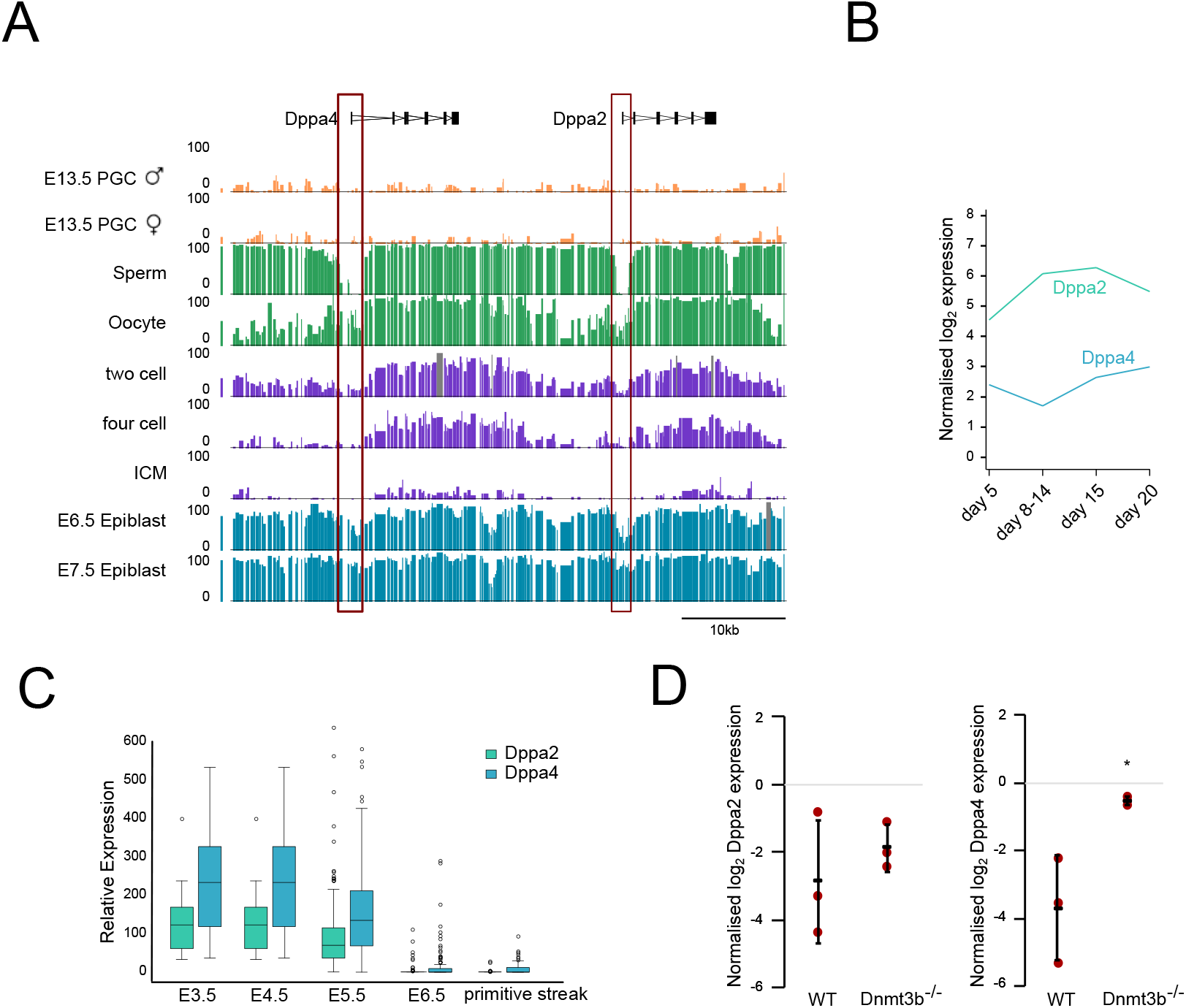
Expression of Dppa2 and Dppa4 coincides with promoter DNA hypomethylation. (A) Whole genome bisulfite data showing percentage DNA methylation across the Dppa2/4 locus in male and female PGCs (orange), sperm and oocytes (green), two cell, four cell embryos and inner cell mass (ICM) of blastocysts (purple) and E6.5 and E7.5 epiblast (blue). Gene structures are shown above tracks and approximate position of DMRs outlined in red boxes. Data reanalyzed from (Wang et al. 2014). (B) Expression levels of Dppa2 (green) and Dppa4 (blue) in non-growing oocytes (postpartum day 5), growing oocytes (postpartum day 8–14 and 15) and fully-grown oocytes (postpartum day 20). Data reanalyzed from (Veselovska et al. 2015). (C) Expression levels of Dppa2 (green) and Dppa4 (blue) in single cells derived from E3.5, E4.5, E5.5, E6.5 embryos and primitive streak. Data reanalyzed from (Mohammed et al. 2017). (D) Expression levels of Dppa2 (left) and Dppa4 (right) in wild type (WT) and Dnmt3b^-/-^ embryos. Bars represent average +/− standard deviation, red dots individual data points * p value<0.05. Data reanalyzed from (Auclair et al. 2014).

### Reducing levels of Dppa2 and Dppa4 leads to a reduction in 2C-like cells and Dux transcription

To test their necessity for ZGA transcripts expression, we first performed Dppa2 and Dppa4 knockdowns in MERVL::tdTomato/ Zscan4c::eGFP reporter serum ESCs (Eckersley-Maslin et al. 2016). Expression of either reporter accurately labels 2C-like cells (Macfarlan et al. 2012; Zalzman et al. 2010; Eckersley-Maslin et al. 2016). Cells were transfected with either control, Dppa2 or Dppa4 targeting siRNAs for 4 days, achieving 92% and 74% knockdown efficiency respectively (Supplemental Figure 4A). Analysis of the MERVL::tdTomato/ Zscan4c::eGFP reporters by flow cytometry revealed a dramatic depletion of the 2C-like ESC population (Figure 4A), which was consistently reflected in the expression of selected ZGA transcripts by qRT-PCR (Supplemental Figure 4B). To further investigate the transcriptional changes occurring after Dppa2 or Dppa4 knockdown, we performed RNA-sequencing. The majority of differentially expressed transcripts were downregulated and overlapped with 2C-like ZGA transcripts (Figure 4B) and were similarly deregulated between Dppa2 and Dppa4 knockdowns (Figure 4C, Supplemental Table 3 and 4). In addition to ZGA transcripts, there was milder downregulation of a second group of genes in the knockdown samples which contained many lineage markers such as the gametogenesis genes Syce1, Sohlh2 and Mael (Figure 4B), consistent with knockout ESC studies (Madan et al. 2009). Thus, while Dppa2 and Dppa4 likely have additional roles, the largest changes in gene expression occurred at the 2C-like ZGA transcripts. The downregulated transcripts were expressed at the time of ZGA in preimplantation embryos (Figure 4D). Transcripts expressed in 2C-like ESCs (Eckersley-Maslin et al. 2016) as well as those that are upregulated following Dppa2, Dppa4 or Dux overexpression (Hendrickson et al. 2017) or following CAF-1 (Ishiuchi et al. 2015) or LINE1 knockdown (Percharde et al. 2018) were all downregulated following Dppa2 or Dppa4 knockdown (Supplemental Figure 4C), indicating the same set of ZGA transcripts is being regulated by Dppa2 and/or Dppa4 irrespective of how they are defined. Importantly, transcript levels of the Dux transcription factor were barely detected following Dppa2 and Dppa4 knockdown (Figure 4E). Therefore, Dppa2 and Dppa4 knockdown results in a decrease in Dux expression, ZGA transcripts and cells in the 2C-like state.

**Figure 4:**
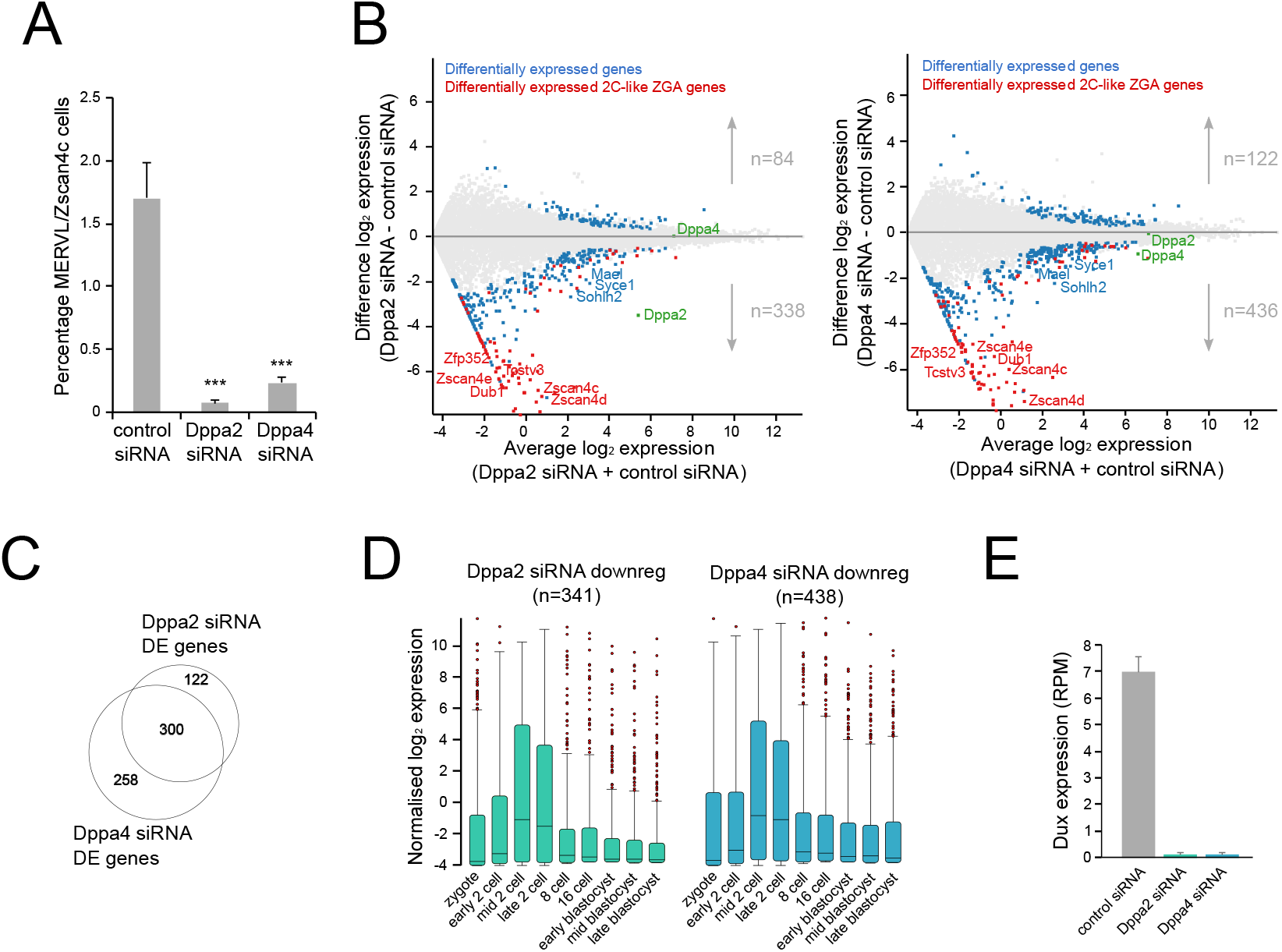
Knockdown of Dppa2 or Dppa4 reduces expression of ZGA transcripts. (A) Flow cytometry analysis of reporter ESCs showing the percentage of 2C-like cells (Zscan4c+ and/or MERVL+) following treatment with either control or target siRNA. Error bars represent standard deviation of 3–6 biological replicates. *** p-value < 0.001 two-tailed equal variance t-test. (B) MA plots showing average log2 expression versus difference in log2 expression for control siRNA and Dppa2 siRNA treated (left) or Dppa4 siRNA treated (right) ESCs, analysed by RNA sequencing. Differentially expressed genes are highlighted in blue, and differentially expressed ZGA transcripts expressed in 2C-like ESCs highlighted in red. Dppa2 and Dppa4 are indicated. (C) Overlap between differentially expressed (DE) genes following Dppa2 or Dppa4 siRNA treatment compared to control siRNA treated cells. (D) Expression pattern of differentially downregulated genes following Dppa2 siRNA treatment (left, green) or Dppa4 siRNA treatment (right, blue) during preimplantation development. Preimplantation data from (Deng et al. 2014). (E) Expression levels of Dux transcript in control siRNA (grey), Dppa2 siRNA (green) and Dppa4 siRNA (blue) treated ESCs.

### Dppa2 and Dppa4 are both necessary for ZGA gene activation

To confirm that Dppa2 and Dppa4 are required for ZGA gene transcription, we generated single and double knockout ESCs deficient for Dppa2 and/or Dppa4 using CRISPR-Cas9 targeting in MERVL::tdTomato reporter cells. Knockout of either or both proteins was confirmed by Western Blotting (Figure 5A). Strikingly, flow cytometry analysis of the MERVL::tdTomato reporter revealed both single and double knockout ESCs completely lacked the 2C-like subpopulation (Figure 5B). Despite the loss of the 2C-like state, the cells retained mRNA and protein expression of the pluripotency markers Nanog and Oct4 (Supplemental Figure 4A-C), grew at a similar rate and were able to be passaged for over 20 generations (data not shown), consistent with previous reports (Madan et al. 2009; Nakamura et al. 2011). Thus, these results indicate that the 2C-like state, as well as Zscan4 protein expression, is dispensable for sustained pluripotency in culture.

**Figure 5:**
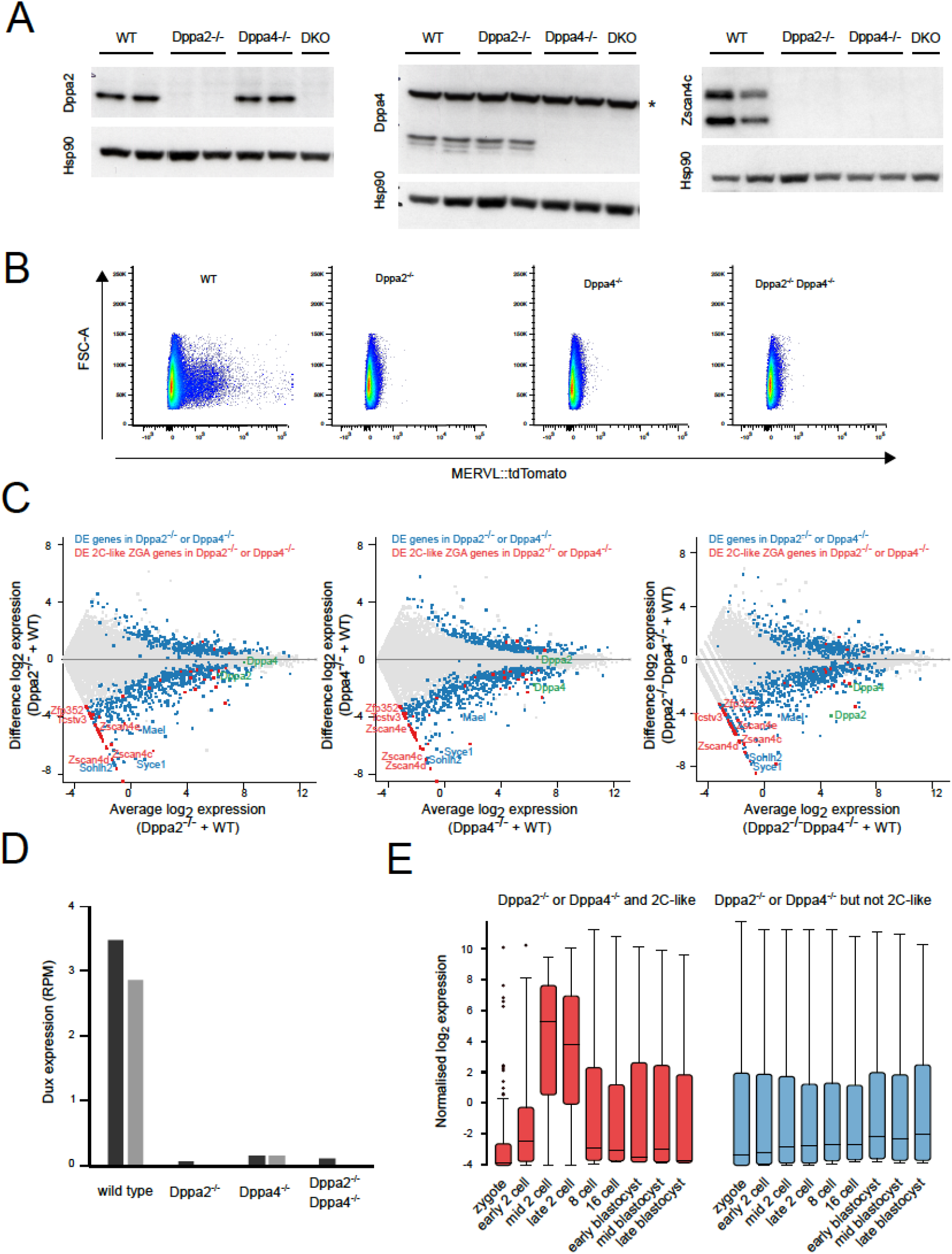
Knockout of Dppa2 and/or Dppa4 in ESCs abolishes 2C-like cells and ZGA transcript expression. (A) Western blotting for Dppa2 (left), Dppa4 (middle) and Zscan4c (right) in individual clones for wild type (WT), Dppa2^-/-^, Dppa4^-/-^ and double Dppa2^-/-^Dppa4^-/-^ (DKO) ESCs. Two clones are shown for WT and single KO and one clone for DKO. Hsp90 is used as loading control. Note the presence of a non-specific band (denoted by *) in the Dppa4 blot. (B) Flow cytometry plots showing expression of MERVL::tdTomato reporter (x-axis) in wild type (WT), Dppa2^-/-^, Dppa4^-/-^ and Dppa2^-/-^Dppa4^-/-^ DKO ESCs. (C) MA plots showing average log2 expression versus difference in log2 expression for wild type (WT), Dppa2^-/-^ (left), Dppa4^-/-^ (middle) or Dppa2^-/-^Dppa4^-/-^ ESCs, analysed by RNA sequencing. Differentially expressed genes in either Dppa2^-/-^ or Dppa4^-/-^ ESCs are highlighted in blue, and differentially expressed ZGA transcripts expressed in 2C-like ESCs highlighted in red. Dppa2 and Dppa4 are indicated. (D) Expression levels of Dux transcript in wild type (WT), Dppa2^-/-^, Dppa4^-/-^ or Dppa2^-/-^Dppa4^-/-^ ESCs. (E) Expression patterns during preimplantation development of differentially expressed genes in either Dppa2^-/-^ or Dppa4^-/-^ ESCs and overlapping (red, left) or not overlapping (blue, right) 2C-like ZGA transcripts. Preimplantation data from (Deng et al. 2014).

Next, we performed RNA sequencing to further investigate the transcriptional changes that occur following loss of Dppa2 and/or Dppa4. We observed a dramatic depletion of 2C-like ZGA transcripts including the Zscan4 cluster, Tcstv3 and Zfp352 (Figure 5C, Supplemental Table 5 and 6), consistent with the knockdown experiments. The absence of Zscan4 cluster transcripts was validated at the protein level by Western Blotting (Figure 5A). Furthermore, expression of the ZGA transcription factor Dux was completely absent in the knockout cells (Figure 5D). The intersection of differentially expressed genes from either Dppa2^-/-^ or Dppa4^-/-^ ESCs were similarly deregulated in the Dppa2^-/-^Dppa4^-/-^ ESCs (Figure 5C) suggesting the three genotypes are largely indistinguishable transcriptionally. These transcriptional changes were validated in independent samples by qRT-PCR (Supplemental Figure 5A). Similar to the knockdown experiments, we also observed a second group of misregulated genes in the knockout ESCs which included many lineage specific genes including Mael, Syce1 and Sohlh2, consistent with previous findings (Madan et al. 2009). However, unlike the differentially expressed 2C-like ZGA transcripts, this second group of differentially expressed genes are not normally upregulated at the time of ZGA during preimplantation development (Figure 5E) and therefore likely represent a separate independent function of Dppa2 and Dppa4 outside of regulating 2C-like ZGA transcription. Therefore, Dppa2 and Dppa4 are necessary for ZGA transcripts expression in ESCs.

Next, we performed rescue experiments in the double knockout ESCs. Consistent with our initial screen, overexpression of Dppa4 and, to a lesser extent, Dppa2 in wild-type (WT) cells upregulated the 2C-like cell fraction by flow cytometry (Figure 6A, Supplemental Figure 6A). Expression of 2C-like ZGA transcripts, including Dux (Figure 6B) was increased. Overexpressing both Dppa2 and Dppa4 resulted in a larger upregulation in 2C-like cells and ZGA transcripts than either alone (Figure 6A-B), consistent with them acting in a complex (Nakamura et al. 2011). Consistently, in Dppa2/Dppa4 null ESCs, Dppa2 was not able to induce the 2C-like state or associated transcripts, and Dppa4 alone led only to a modest increase. Moreover, overexpression of Zscan4c was not able to rescue the Dppa2/Dppa4 knockout phenotype (Supplemental Figure 6B), suggesting that Zscan4c requires Dppa2 and Dppa4 to enhance the 2C-like cell state. Importantly, reintroduction of both Dppa2 and Dppa4 resulted in a substantial increase in the MERVL::tdTomato positive cell fraction (Figure 6A) and associated transcripts, including Dux (Figure 6B). Therefore, Dppa2 and Dppa4 together drive the 2C-like cell state and ZGA transcriptional network.

**Figure 6:**
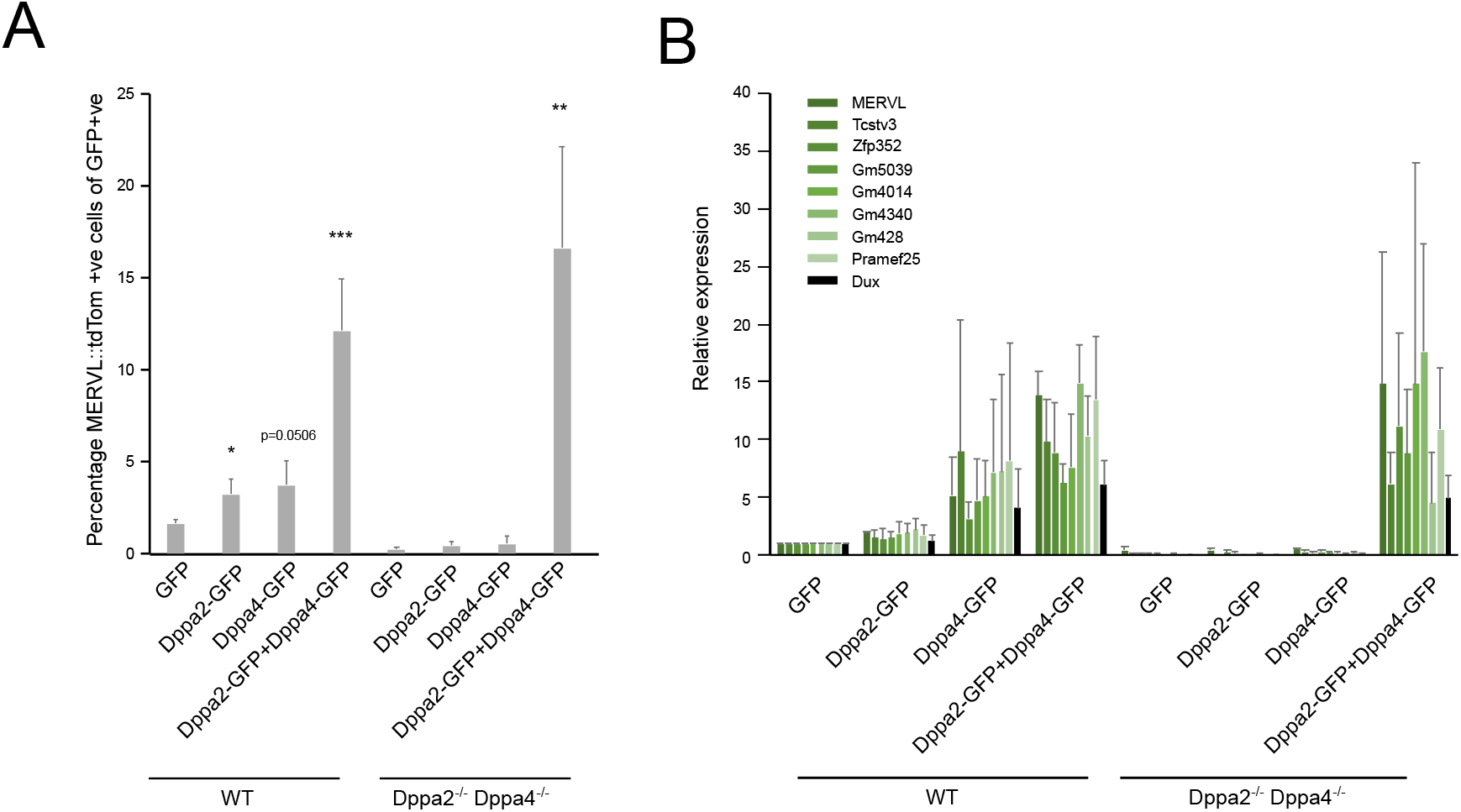
Rescue of Dppa2 and Dppa4 restores 2C-like cells and ZGA transcript expression. Rescue experiments in wild type (WT, left) and Dppa2/4 DKO (right) ESCs. Cells were transfected with GFP, Dppa2-GFP, Dppa4-GFP or Dppa2-GFP with Dppa4-GFP constructs for 48 hours. (A) Expression of MERVL::tdTomato reporter measured by flow cytometry (B) Expression of ZGA transcripts measured by quantitative RT-PCR. Differences are statistically significant (homoscedastic two-tailed t-test, * p-value μ0.5, ** p-value μ0.01, *** p-value < 0.001, **** p-value < 0.0001). Error bars represent average plus standard deviation of three biological replicates.

### Dppa2 and Dppa4 directly bind and regulate the transcription factor Dux

Our results so far have revealed a role for Dppa2 and Dppa4 in regulating the 2C-like state and ZGA transcripts. Additionally, Dppa2 and Dppa4 are necessary and sufficient to regulate expression of the ZGA transcription factor Dux, which itself has recently been shown to regulate a similar ZGA transcriptional programme (De Iaco et al. 2017; Hendrickson et al. 2017; Whiddon et al. 2017). To determine whether Dppa2 and Dppa4 act to regulate Dux directly or whether they exert their effects through parallel pathways, we determined whether Dux is required for Dppa2 and Dppa4 to regulate the ZGA transcripts. Wild type (WT) and Dux knockout ESCs (De Iaco et al. 2017) were cultured in serum conditions and transfected with constructs containing Dppa2, Dppa4 or both Dppa2 and Dppa4 constructs simultaneously, and compared to those receiving an empty vector (Supplemental Figure 7A-B). While Dppa2 and/or Dppa4 were able to induce expression of the ZGA transcripts in WT cells, this ability was abolished in Dux knockout cells (Figure 7A). Therefore, Dux is required for the transcriptional effects exerted by Dppa2 and Dppa4.

**Figure 7:**
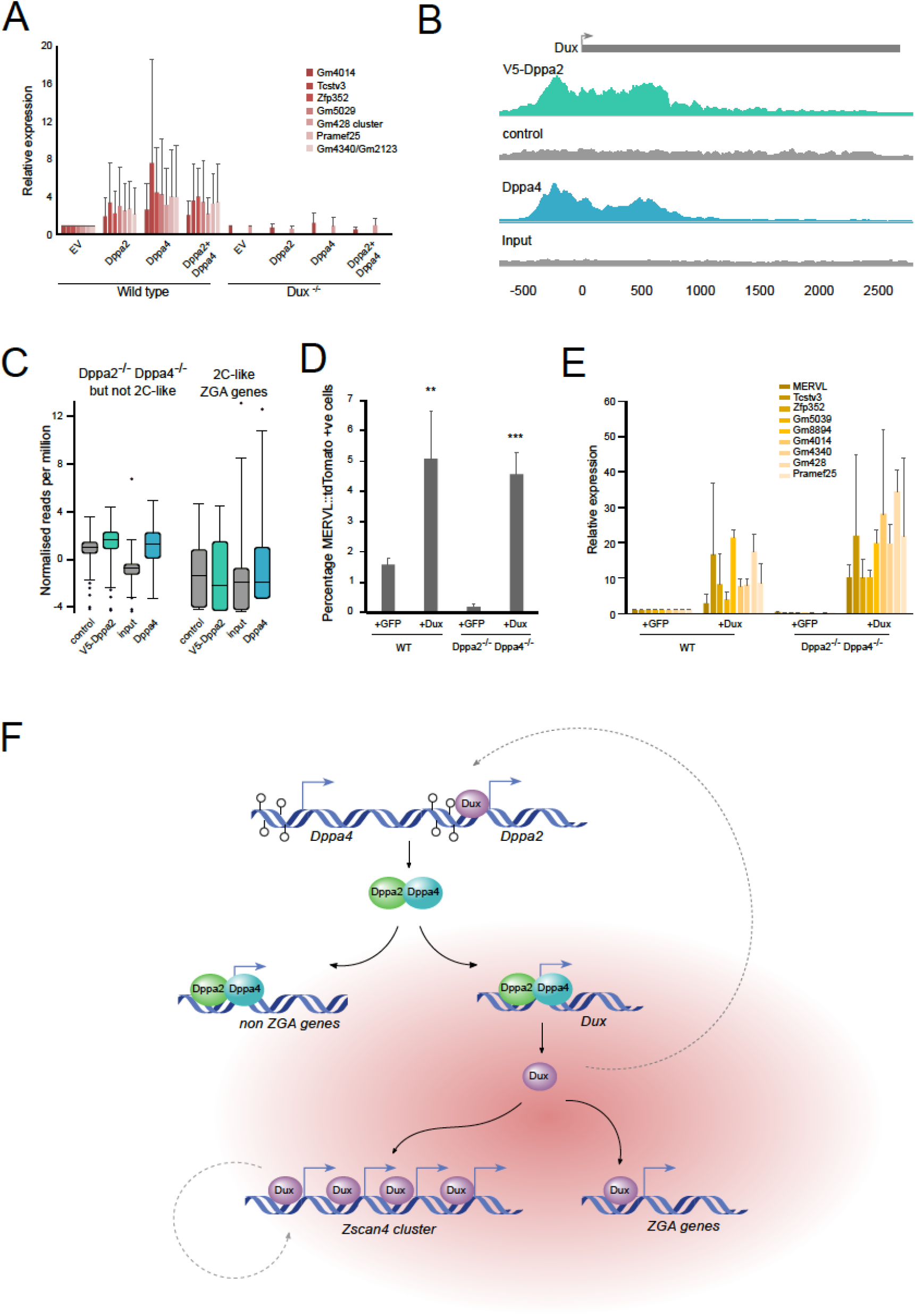
Dppa2 and Dppa4 bind and regulate Dux which in turn is required to upregulate ZGA transcripts. (A) Quantitative RT-PCR analysis of 2C-like ZGA transcripts following transient transfection of untagged Dppa2 and/or Dppa4 in wild type (left) and Dux^-/-^ (right) ESCs, using transfection of an empty vector (EV) as a control. (B) ChIP-seq analysis of Dppa2-V5 (top, green) and endogenous Dppa4 (bottom, blue) binding to Dux consensus sequence and promoter region. Dux transcriptional unit is denoted in grey, scale represents base pairs relative to TSS. Control ESCs not expressing Dppa2-V5 (second row, grey) or input (bottom row, grey) are shown. Dppa2 data reanalyzed from (Engelen et al. 2015), Dppa4 data reanalyzed from (Hernandez et al. 2018). (C) Box whisker plots showing enrichment of Dppa2 (green) and Dppa4 (blue) at differentially expressed genes following Dppa2 or Dppa4 knockdown which either do not (left) or do (right) overlap with 2C-like ZGA transcripts. Probes were made across transcriptional start sites (+/− 500bp) and normalized counts per million reads determined and compared to control/input (grey). (D) Flow cytometry analysis of reporter ESCs showing the percentage of MERVL::tdTomato+ve 2C-like cells in WT or Dppa2^-/-^Dppa4^-/-^ ESCs transfected with Dux or GFP positive sorted cells as a control. Error bars represent standard deviation of 3biological replicates. Differences are statistically significant (** p-value < 0.01 *** p-value < 0.001 two-tailed homoscedastic t-test). (E,F) Quantitative RT-PCR analysis of (E) 2C-like ZGA transcripts in cells in WT or Dppa2-/Dppa4^-/-^ ESCs transfected with Dux or GFP control. (F) Promoter DNA demethylation (open circles) enables expression of Dppa2 (green) and Dppa4 (blue), which bind to both non ZGA genes and, under permissive conditions (red cloud), the ZGA transcription factor Dux, inducing its expression. Dux (purple) then binds and activates ZGA genes including the Zscan4 cluster of genes. Dotted grey lines represent positive feedback loops.

Next, we analysed published Dppa2 and Dppa4 ChIP-seq data (Engelen et al. 2015; Klein et al. 2018) to determine whether these proteins directly bind Dux. Due to the repetitive nature of the Dux transcript, we built a custom chromosome containing the Dux consensus sequence along with upstream genomic sequence. Importantly, we observed clear enrichment of Dppa2 and Dppa4 binding across the promoter and into the Dux transcript itself in E14 ESCs (Figure 7B). Furthermore, Dppa4 similarly bound Dux in P19 embryonal carcinoma cells but not in 3T3 fibroblasts (Supplemental Figure 7C). Thus, Dppa2 and Dppa4 directly bind to Dux in pluripotent stem cells, consistent with when Dux is expressed. We next investigated whether Dppa2 and Dppa4 bind to other genes that are differentially in Dppa2^-/-^ or Dppa4^-/-^ ESCs. There was a strong enrichment for both Dppa2 and Dppa4 at the promoters of the non 2C-like ZGA genes (Figure 7C), including Syce1, Sohlh2 and Mael (Supplemental Figure 7D-F), again confirming a separate role for Dppa2 and Dppa4 in regulating non ZGA transcripts in ESCs. Importantly, Dppa2 and Dppa4 did not bind to the transcriptional start site of other 2C-like ZGA transcripts (Figure 7C), including the Zscan4 cluster, Gm428 and Dub1 (Supplemental Figure 7G-I), which are direct targets of Dux itself (Hendrickson et al. 2017). Lastly, we overexpressed Dux in the Dppa2^-/-^Dppa4^-/-^ ESCs (Supplemental Figure 7J). Our results support a model by which Dppa2 and Dppa4 act by directly regulating levels of the Dux transcription factor which in turns acts to bind and promote expression of a zygotic transcriptional programme in ESCs. Therefore, expressing Dux in the absence of Dppa2 and Dppa4 should restore the 2C-like state and associated ZGA transcripts. Indeed flow cytometry analysis of the MERVL::tdTomato reporter revealed that Dux overexpression is able to induce the 2C-like state in both WT and Dppa2^-/-^Dppa4^-/-^ ESCs (Figure 7D). Furthermore, expression of 2C-like ZGA transcripts were induced following Dux overexpression (Figure 7E). In summary, Dppa2 and Dppa4 induce the 2C-like state and associated ZGA transcripts by directly binding and activating the ZGA transcription factor Dux which is then able to bind and activate downstream 2C-like ZGA target genes.

## Discussion

Initiation of transcription of the zygotic genome is a critical step in embryogenesis. To understand its molecular regulation, we performed a screen for positive chromatin and epigenetic regulators of ZGA transcription using 2C-like ESCs as a model, identifying, amongst others, Dppa2 and Dppa4 as activators of ZGA gene transcription in ESCs. While there are many known repressors of the 2C-like state to date, only the transcription factor Dux has been identified as a positive regulator of ZGA transcription (De Iaco et al. 2017; Hendrickson et al. 2017; Whiddon et al. 2017). However, transcription of Dux itself is not initiated until the minor wave of ZGA and its regulation remains unknown. Here, we propose a model in which promoter DNA demethylation during global epigenetic reprogramming enables expression of Dppa2 and Dppa4 in the germ line and oocytes. Dppa2 and Dppa4 then directly bind and upregulate both non ZGA genes as well as, under permissive conditions, the ZGA transcription factor *Dux* (Figure 7F). Dux is subsequently able to bind and activate downstream ZGA transcription including the *Zscan4* cluster. Several feedback loops reinforce this system, including Zscan4 induced stabilization, as well as Dux induced upregulation of *Dppa2*. Our study provides crucial insights into the molecular hierarchy that triggers ZGA transcription, and links it with epigenetic reprogramming in the germline.

The existence of a 2C-like state in ESCs represents a useful *in vitro* model system for studying ZGA, making many molecular and screening-based studies possible. This state is characterised by an increase in chromatin mobility (Boskovic et al. 2014), decondensed chromocenters (Akiyama et al. 2015; Ishiuchi et al. 2015), and increased chromatin accessibility (Eckersley-Maslin et al. 2016). Consistently, depletion of factors involved in chromatin assembly (Ishiuchi et al. 2015) or treatment with inhibitors which ultimately induce chromatin decompaction (Dan et al. 2015; Macfarlan et al. 2012) increases the proportion of these cells in culture. While 2C-like cells exhibit lower levels of DNA methylation, this hypomethylation occurs after the cells enter the state (Eckersley-Maslin et al. 2016; Dan et al. 2017), and consequently, neither DNA hypomethylation nor deletion of DNA methyltransferases induce ZGA transcription in ESCs (Eckersley-Maslin et al. 2016). Instead, the histone modifications may be important for repressing ZGA genes, as knockdown or knockout of many repressive epigenetic regulators, including the histone demethylase Kdm1a (Macfarlan et al. 2011), histone methyltransferases Ehmt2 (Macfarlan et al. 2012), heterochromatin protein HP1 (Maksakova et al. 2013), components of the PRC1.6 subcomplex (Rodriguez-Terrones et al. 2017), amongst others (Fujii et al. 2015; Dan et al. 2014; Maksakova et al. 2013; Rodriguez-Terrones et al. 2017), have also been shown to repress the 2C-like state.

Any positive regulator of a transcriptional programme must be present prior to these genes being expressed. In this way, Dux, while a master regulator of all downstream ZGA genes, cannot be the initial trigger for ZGA as it is itself transcribed in the minor wave of ZGA. In primordial germ cells, the promoters of *Dppa2* and *Dppa4* are demethylated, coinciding with their transcription. They remain robustly expressed through preimplantation development, including at the time of ZGA, until the onset of gastrulation when expression rapidly ceases and their promoters acquire DNA methylation. Dppa2 is highly expressed in oocytes and zygotes prior to ZGA, and while Dppa4 transcripts are less abundant, proteins for both Dppa2 and Dppa4 are readily detectable (Pfeiffer et al. 2011). Additionally, Dppa2 and Dppa4 are expressed at the time during iPSC reprogramming when 2C-like ZGA transcripts are transiently expressed. Furthermore, Dppa2 and Dppa4 are homogeneously expressed across all ESCs, not just 2C-like cells (data reanalyzed from (Eckersley-Maslin et al. 2016)). However, this raises the question as to why the ZGA genes are expressed in only a small subset of and not all ESCs. Activation of the 2C-like state is likely a multifaceted process requiring not only presence of the upstream activators Dppa2 and Dppa4, but also chromatin decompaction and/or reduced expression of repressors such as Kap1 or PRC1 and their modifications. Once activated, factors such as Zscan4c may act to stabilize and prolong expression of the transcriptional programme which then requires repressors, such as LINE-1/Nucleolin complex (Percharde et al. 2018), NuRD or Caf-1(Campbell et al. 2018; Ishiuchi et al. 2015), to repress it once again. In this way Dppa2 and Dppa4 regulate the entry into the 2C-like state during permissive conditions by directly activating the ZGA major transcription factor Dux, which subsequently activates the remainder of the ZGA transcripts in the cell.

Dppa2/4 knockout ESCs retained expression of pluripotency markers and could be passaged for over 20 generations, consistent with previous reports (Nakamura et al. 2011; Madan et al. 2009). Therefore, while Dppa2 and Dppa4 have no major effect on ESC pluripotency, their absence completely abolishes 2C-like cells, indicating that the 2C-like state is dispensable for ESC pluripotency. This is in contrast with studies reporting that the ZGA marker Zscan4 is required for long-term ESC culture maintenance and telomere stability (Zalzman et al. 2010) as Dppa2 and/or Dppa4 knockout completely abolishes Zscan4 expression.

In this study, we used Zscan4c as a positive control in the candidate based screen. Zscan4c is one of a tandemly encoded family of zinc finger and SCAN domain containing proteins which are expressed in early embryos and in 1–5% of ESCs (Zalzman et al. 2010), including the rarer MERVL positive 2C-like cells (Eckersley-Maslin et al. 2016; Rodriguez-Terrones et al. 2017). Zscan4c has been implicated as a positive regulator of 2C-like cells (Hirata et al. 2012; Amano et al. 2013), which we have confirmed here. However, Zscan4c is unable to induce the 2C-like state or ZGA transcription in the absence of Dppa2, Dppa4 or Dux (data not shown). While it remains to be determined whether Zscan4c or other members of the Zscan4 cluster are necessary for ZGA transcription, our results suggest that Zscan4c may act to stabilize or reinforce the 2C-like state rather than to induce it. Interestingly, as well as Zscan4c, Sp110 is also upregulated in 2C-like cells and following Dux overexpression and was also identified in our screen as a positive regulator of the 2C-like state. It will be interesting to see if it may also act as a reinforcer of the 2C-like state in ESCs.

After Zscan4c, Dppa4 was the strongest identified inducer of ZGA transcription and was therefore selected, along with the closely related Dppa2, for further study. Interestingly, the family member Dppa3 (also known as Stella or Pgc7) was not able to induce the 2C-like state, consistent with a previous report (Huang et al. 2017). Dppa2 and Dppa4 physically interact and bind to euchromatin in pluripotent stem cells (Nakamura et al. 2011; Klein et al. 2018). Here, we show that Dppa2 and Dppa4 are regulated by DNA methylation and are both necessary to induce an early embryonic transcriptional network by directly binding and regulating the ZGA transcription factor Dux in ESCs. *In vivo*, Dux knockdown results in impaired early embryonic development and defective ZGA (De Iaco et al. 2017). Importantly, injection of a dominant negative form of Dppa2 lacking the SAP domain into zygotes induces two cell arrest (Hu et al. 2010), suggesting that our results showing Dppa2 and Dppa4 regulate the ZGA transcriptional network in ESCs may also occur in embryos.

In summary, in this study we performed a candidate-based screen to identify epigenetic and chromatin regulators of the 2C-like state and ZGA transcriptional programme. Amongst these were Dppa2 and Dppa4 which act together to bind and upregulate the ZGA transcription factor Dux, amongst other non-ZGA targets. Depleting Dppa2 and Dppa4 levels reduces the 2C-like population and ZGA transcription which can be restored by reintroducing either both Dppa2 and Dppa4 together, or the downstream factor Dux. In conclusion, our findings shed important insights into the molecular mechanisms regulating ZGA transcription.

## Materials And Methods

### Gateway cloning

Sequence verified cDNA sequences lacking stop codons were amplified from plasmids purchased from ThermoFisher using primers containing AttB1 and AttB2 sequences and cloned into pDONR221 vector. Gateway cloning was then used to transfer the cDNA sequences into an in-house built pDEST vector containing a CAG promoter and an in-frame C-terminal eGFP coding sequence and blasticidin resistance by IRES fusion. Expression plasmids were sequence verified by Sanger Sequencing prior to use and are available upon request.

### Cell culture

E14 mouse embryonic stem cells were grown under standard serum/LIF conditions (DMEM 4,500 mg/l glucose, 4 mM L-glutamine, 110 mg/l sodium pyruvate, 15% fetal bovine serum, 1 U/ml penicillin, 1 mg/ml streptomycin, 0.1 mM nonessential amino acids, 50 mM b-mercaptoethanol, and 10^3^ U/ml LIF). Single MERVL::tdTomato and double MERVL::tdTomato/Zscan4c::eGFP reporter cell lines are described in (Eckersley-Maslin et al. 2016) and Dux knockout cells described in (De Iaco et al. 2017). Transfections were performed using lipofectamine on pre-plated cells in 6 well or 10cm plate formats. Flow cytometry analysis was performed using BD LSR Fortessa and sorts performed on BD Aria III or BD Influx High-Speed Cell Sorter. siRNA transfections were performed by transfecting Dharmacon siRNA ON-TARGETplus siRNA SMARTpool at a final concentration of 50nM with lipofectamine.

### Generation of Dppa2 and Dppa4 CRISPR knockout ESCs

CRISPR knockout ESCs were performed as previously described (Ran et al. 2013). Guide RNAs were designed against exons 2 and 3 of both Dppa2 and Dppa4 using CRISPR design (crispr.mit.edu). Cells were transfected with a single guide targeting Dppa2 and/or Dppa4, FACS sorted following 24 hours into single cells, and clones screened by surveyor assay and genomic DNA PCR. Successfully targeted clones were validated by Western Blotting. Dppa2 single knockout clones (clone 5 and clone 12) used in this study were generated using a guide RNA targeting Dppa2 exon 2 (ACCTTAGACCACACACCACCAGG); Dppa4 single knockout clones (clone 23 and clone 29) with a guide RNA targeting Dppa4 exon 2 (CTGCAAAGGCTAAAGCAACGGGG); Dppa2/Dppa4 double knockout clone (clone 43) was generated using a guide targeting Dppa2 exon 3 (TAACTTGAGTACGGATGGCAAGG) together with the guide RNA targeting Dppa4 (as above).

### RNA isolation, qPCR and RNA-sequencing

RNA was isolated using Qiagen RNA-DNA allprep columns or TriReagent (Sigma) and treated with DNAseI (Ambion DNA-free DNA Cat #1311027 or ThermoFisher RNase-free EN0525) using manufacturer’s instructions. cDNA was generated using 0.5–1 μg RNA (Thermo RevertAid #K1622), and quantitative RT-PCR performed using Brilliant III SYBR master mix (Agilent Technologies #600882). Relative quantification was performed using the comparative CT method with normalisation to CycloB1 levels. Primer sequences are available upon request. Opposite strand-specific polyA RNA libraries were made using 1 μg DNAse treated RNA at the Sanger Institute Illumina bespoke pipeline and sequenced as single end 50bp reads using the Illumina HiSeq 2500 Rapid Run platform.

### Candidate screen selection

Candidate 2C-like ZGA regulators were selected based on the following criteria. First, genes expressed in oocytes/zygotes/two cell embryos (RPKM>1 in (Deng et al. 2014) data) were selected and further filtered based on gene ontology (AmiGO Ontology search for ‘chromatin’ associated genes). This gave a total list of 84 candidate genes which were manually curated to remove non nuclear proteins and overlapping transcripts. Genes were finally filtered to include those which had sequence verified cDNA clones commercially available.

### Data Analysis

Raw FastQ data were trimmed with Trim Galore (v0.4.3, default parameters) and mapped to the mouse GRCm38 genome assembly using Hisat2 v2.0.5. Data were quantitated at mRNA level using the RNA-seq quantitation pipeline in SeqMonk software (http://www.bioinformatics.babraham.ac.uk/projects/seqmonk/). Strand specific quantification was performed using mRNA probes and cumulative distributions matched across samples. Differentially expressed genes were determined using DESeq2 (p-value 0.05, with multiple testing correction) and Intensity difference filter (p-value 0.05), with the high-confidence DE genes defined as the intersection between the two statistical tests. For Dux analysis, fastq files for RNA-seq and published ChIP-seq data were realigned to the mouse dux repeat (AM398147.1) and read counts normalised to total read counts. Graphing and statistics was performed using Seqmonk, Excel or RStudio.

### Western Blotting

Western Blotting was performed using 50 μg protein extracted using detergent buffer (10mM Tris-HCl, pH 7.4, 150mM NaCl, 10mM KCl, 0.5% NP-40) containing a protease inhibitor cocktail (Sigma, P2714) and quantified by Bio-Rad Protein Assay Dye Reagent. Proteins were resolved using 4–12% SDS-PAGE gels (Expedon, NBT41212) and blotted on PVDF membranes. Following blocking in 5% skim milk/0.01% Tween/PBS, membranes were incubated with primary antibodies for 3 hours to overnight. Secondary Horseradish peroxidase-conjugated secondary antibodies (Santa Cruz 1:3000) were incubated for 1 hour and detection carried out with enhanced chemiluminescence (ECL) reaction (GE Healthcare, RPN2209). Primary antibodies: anti-Zscan4 (Millipore, 2793611, 1:500), anti-Dppa2 (Millipore MAB4356, 1:500), Dppa4 (Santa Cruz sc-74614, 1:200) and anti-HSP90 (Abcam, ab13492, 1:5000).

## Acknowledgments

The authors thank all members of the Reik laboratory for helpful discussions and Aled Parry for reading the manuscript. We also thank Simon Andrews and Felix Krueger for bioinformatics support, Nathalie Smerdon for assistance with high-throughput sequencing and Rachael Walker for assistance with flow cytometry. We also thank Didier Trono (EPFL) for kindly providing Dux knockout ESCs and Dux constructs. M.A.E-M was supported by an EMBO Fellowship (ALTF938–2014) and Marie Sklodowska-Curie Individual Fellowship. Research in the Reik lab is supported by BBSRC (BB/K010867/1), Wellcome Trust (095645/Z/11/Z) and EU EpiGeneSys.

## Author Contributions

M.A.E.-M. and W.R. conceived and designed the study. M.A.E.-M. performed and analysed experiments, performed bioinformatic analysis and wrote the paper. C.A.-C. performed Western Blotting and Dux knockout experiments. M.B. assisted with screen and performed Dppa2/Dppa4 knockout rescue experiments. E.K. assisted with siRNA knockdown experiments. C.K. performed Dux repeat mapping. M.A.E.-M. and W.R. supervised the study.

